# The Laminar Organization of a Decision Circuit in Orbitofrontal Cortex is Agnostic to the Variable Encoding Scheme

**DOI:** 10.64898/2026.06.01.729414

**Authors:** Alessandro Livi, Manning Zhang, Camillo Padoa-Schioppa, Timothy E. Holy

## Abstract

Economic choices are believed to depend on the orbitofrontal cortex (OFC). Work in primates and rodents indicates that neurons in OFC participate in computing and comparing subjective values, suggesting that different groups of cells constitute the building blocks of a decision circuit. In a recent study (Livi et al., 2025), we examined the laminar organization of this circuit in mice. We found that different decision variables were differentially represented in layer 2/3 (L2/3) and layer 5 (L5). Furthermore, the temporal dynamics of decision signals indicated a combination of feed-forward and feed-back across layers, and pointed to L5 as the locus for winner-take-all value comparison. Importantly, these results were obtained under the constraint that each neuron encoded a single variable. Here, we tested whether our results on laminar organization depended on the categorical framework. We applied LASSO regression to identify a minimal set of variables explaining each neuron’s activity. Even with approximately half of all neurons representing two or more variables, the layer specificity of decision variables was preserved. In addition, Granger Causality Analysis and activity profiles reached similar conclusions as for analyses conducted under the single-variable constraint. We conclude that the decision circuit in OFC exhibits a laminar architecture, independently of whether the representation of decision variables in this area is categorical or mixed.

## Introduction

Neural computation is constrained by cellular properties and patterns of connectivity, and distinct collections of neurons frequently encode different task-related variables. In some cases, cellular and connectivity differences align with anatomical structure, either across brain areas or across layers within a cortical circuit. Elucidating how task variables are represented in neural activity, and how these representations map onto specific cells and circuit architectures, is a longstanding goal of neuroscience.

The structure of neural computation becomes particularly salient in tasks that can be framed in terms of a small number of interpretable variables. In economic choice, extensive experimental and theoretical work has identified within orbitofrontal cortex (OFC) a limited set of variables related to the representation and comparison of offer values and choice outcomes (Padoa-Schioppa and Assad, 2006; Pastor-Bernier et al., 2019; Gore et al., 2023). Using single-unit recordings, some studies have reported that individual neurons encode these variables categorically, such that neural responses are selectively tuned to quantities like the value of a specific offer, the value of the chosen option, or spatial features associated with reward collection, in both rats (Hirokawa et al., 2019) and monkeys (Onken et al., 2019). Other studies have emphasized mixed representations (Rigotti et al., 2013; Kaufman et al., 2022; Tye et al., 2024), with individual neurons encoding multiple variables and little evidence for clustering along specific representational axes (Blanchard et al., 2018; Posani et al., 2025). Most recently, a study revealed an heterogeneous encoding structure where most neurons display mixed selectivity and population encoding is categorical for certain variables, but non-categorical for others (Crimmins et al., 2025).

Despite this substantial body of work, much less is known about how representations of decision variables relate to circuit architecture, including laminar organization and patterns of functional interaction. This gap limits mechanistic understanding of how economic decisions are implemented in cortical networks. For this reason, in a recent study we recorded from neurons in the mouse OFC using two-photon microscopy (Zhang et al., 2024; Livi et al., 2025). Neurons in the mouse OFC encode a variety of decision variables, partly overlapping with those described in primates.

Furthermore, neurons with similar tuning are spatially clustered (Livi et al., 2025). Most notably, these representations were organized across cortical layers: activity in layer 2/3 (L2/3) and layer 5 (L5) differed systematically in the variables they represented and in their temporal profiles, consistent with a laminar structure underlying value-based decision-making (Livi et al., 2025). These results were obtained from an analysis in which each neuron was permitted to encode at most one variable out of a predefined list. Encoding of decision variables in OFC has traditionally been described as categorical (Onken et al., 2019). However, a subsequent study by Crimmins and colleagues (Crimmins et al., 2025) reported that OFC neurons can exhibit mixed selectivity to economic variables. In light of this ongoing debate, it is natural to ask whether the results of our previous work (Livi et al., 2025) arise as a consequence of enforcing categorical encoding, or whether they remain valid when neurons are allowed to encode more than one decision variable.

Here, we address this question by revisiting the laminar organization of OFC decision-related activity using LASSO regression (Tibshirani, 1996; Ranstam and Cook, 2018), a sparse modeling approach that allows neuronal responses to be explained by combinations of task variables without enforcing categorical assignment. This framework provides a principled way to assess whether laminar differences in representational content and temporal dynamics persist when neurons are permitted to encode more than one variable. Using LASSO-derived representations, we examine how decision variables are distributed across layers, how these representations evolve over time, and how they relate to patterns of functional interaction within the circuit. Our findings show that the laminar and temporal organization of decision-related activity in OFC is preserved under this more general representational framework. Together, these results support a robust, layer-specific and categorical independent organization of value-based decision processes in OFC.

## Material and Methods

The present study provides new analyses of previously published imaging and behavioral datasets (Zhang et al., 2024; Livi et al., 2025). All experimental procedures conformed to the NIH Guide for the Care and Use of Laboratory Animals and were approved by the Institutional Animal Care and Use Committee (IACUC) at Washington University in St. Louis. Detailed procedures for surgery, viral injection, behavioral training, neural recordings, image processing, image registration, ΔF/F extractions and initial data curation can be found in Livi et al. (2025). Here, we summarize the components relevant to the new analyses and describe methodological choices that deviate from or extend those earlier studies. Of note, our dataset included neurons imaged from L2/3 and L5. In LASSO analysis and Granger Causality Analysis (GCA) we pooled neurons from different layers; for the remaining analyses, we split layers.

### Animal subjects

Data were collected from 14 mice (7 males, 7 females; animals labeled with prefix “M”; see Supplementary Table 1 in Livi et al. (2025)). After surgical preparation, mice were housed individually and tested during the light phase of a 12 h light/dark cycle. Animals were kept under water restriction and obtained most or all of their daily water intake during behavioral sessions. Body weight was monitored daily; if fluid intake dropped below 0.6 mL or body weight fell below 75% of pre-restriction baseline, supplemental water was provided.

### Choice task

#### Trial structure

The behavioral apparatus was custom-built and controlled using LabView. Mice were head-fixed inside a plastic tube (**Fig.1**). Two odor ports (left/right) delivered olfactory stimuli signaling two juice offers, and two liquid spouts near the mouth detected licks and delivered reward. Each trial began with a vacuum sweep to remove residual odor. Two olfactory stimuli – defined by odor identity (juice flavor) and concentration (juice quantity) – were presented simultaneously from left and right ports for 2.4 – 2.8 s (constant within animal). A brief “go” cue marked the onset of the choice period. The mouse indicated its choice by licking one of the two spouts. Early licks were ignored; trials with no lick within 5 s were aborted. In forced-choice trials, licking the wrong spout yielded a white-noise error signal followed by a repeat of the same offer. Animals choose between their preferred juice (A) and the non-preferred juice (B). Quantities varied over 4 – 5 levels (0 – 7 drops). Spatial arrangement (A left/right) varied pseudo-randomly. Rewards were delivered at ∼150 ms per drop. Delivery duration was fixed within each session to the maximum reward quantity delivered in that session. For example, if the maximum drop quantity in a given session was 7 (independent of juice type), the interval between reward delivery and the onset of the subsequent inter-trial interval (ITI) was fixed at 1050 ms (150 ms × 7) for all trials. This procedure prevented trial-by-trial temporal discounting.

**Figure 1.**
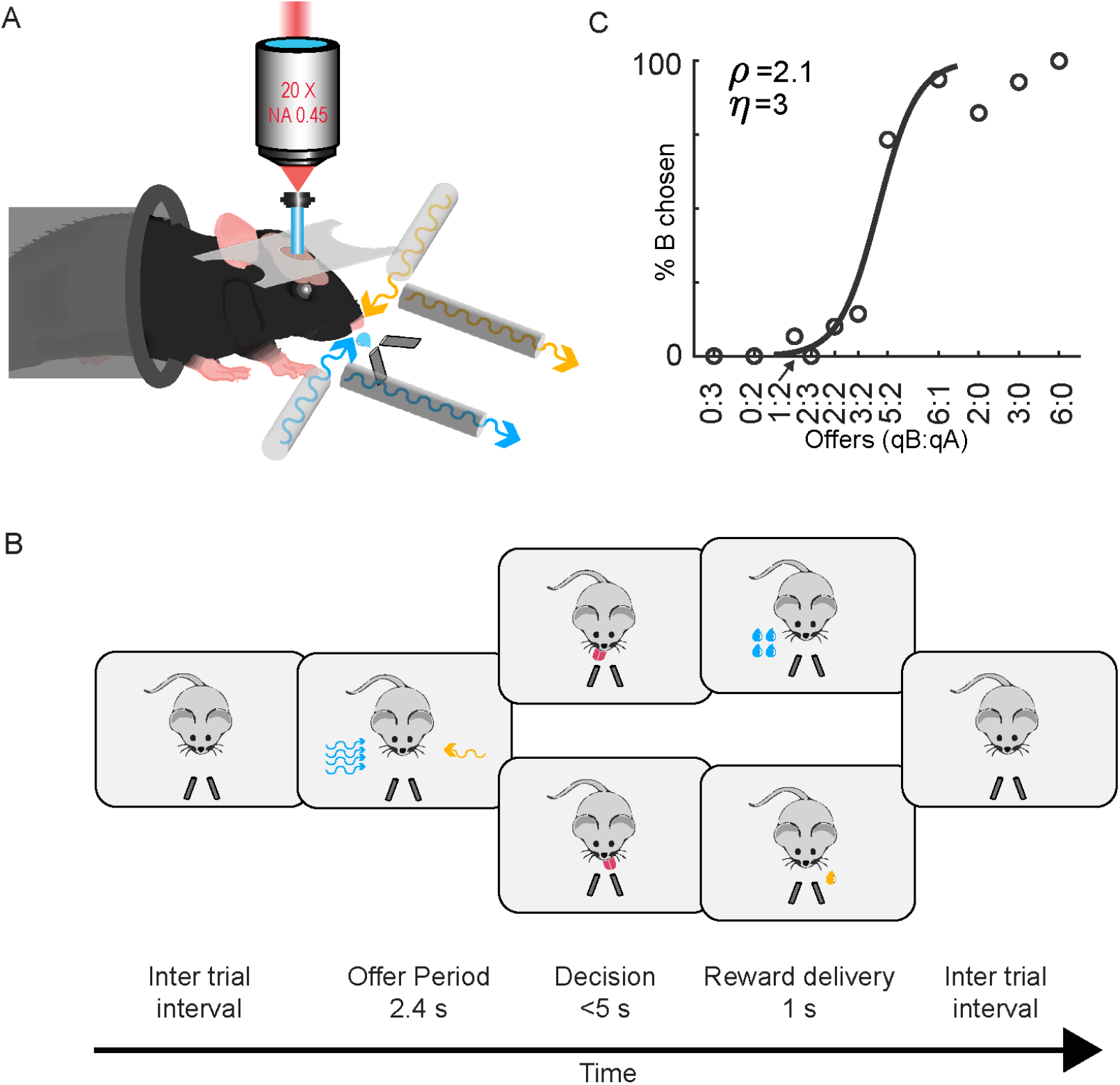
Choice task and neural recordings in OFC. **A**. Apparatus. Mice were head-fixed, restrained inside a tube, implanted with a GRIN lens targeting OFC, and positioned under a two-photon microscope. Olfactory stimuli were delivered through two odor ports (left/right), and two additional ports vacuumed residual odors before each trial. Two reward spouts (left/right) detected the animal’s choice (first lick after the go signal) and delivered the chosen juice. **B**. Choice task. After the intertrial interval (ITI), the animal was presented with two odors whose identities indicated juice types and whose concentrations encoded offered quantities. Following the offer period (2.4 – 2.8 s), a tone signaled the go cue. The mouse had up to 5 s to lick one of the two spouts; reward was delivered immediately afterward. The spatial configuration of offers (A/B left – right) varied pseudo-randomly across trials. **C**. Example session. Behavioral choices are plotted as the percentage of trials in which the animal chose juice B (y-axis) versus the log quantity ratio log(qB/qA) (x-axis). The logistic fit (black curve) provided the relative value (ρ) and choice accuracy (η). Only behavioral sessions satisfying predefined criteria (**Methods**) were used for neural analyses.

#### Odor Stimuli

Two monomolecular odors, octanal (#0568, Sigma) and octanol (#472328, Sigma), were used because they are volatile, easily discriminated (Laska et al., 2008), and typically neutral for mice (Saraiva et al., 2016). Odor-juice associations were pseudo-randomized across animals. The odor concentration increased monotonically with quantity on a log_2_ scale (1, 2, 4, 8 ppm → quantities 1-4).

### Analysis of choice data

All analyses were performed in Matlab 2021b (MathWorks). Behavioral choices were analyzed using logistic regression (Padoa-Schioppa, 2022). First, we fit a full model incorporating potential side bias that predicts the outcome of each trial:

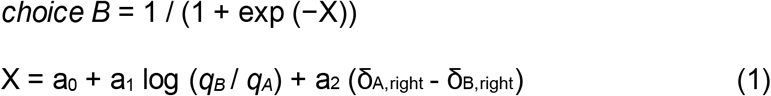

where *choice B* = 1 if the animal chose juice B and 0 otherwise, *q*_*A*_ and *q*_*B*_ are the offered quantities for juices A and B, and δ_J,right_ = 1 if juice J was offered on the right and 0 otherwise (J = A or B). For each session, the logistic regression (maximum likelihood) returned two sigmoid functions with the same steepness and different flex points, one for A offered on the right the other for A offered on the left. From fitted parameters a_0_, a_1_ and a_2_, we computed the choice accuracy *η* = a_1_, the (subjective) relative value of juices *ρ* = exp(−a_0_/a_1_), and the side bias *ε* = −a_2_/a_1_. Intuitively, *η* (also termed inverse temperature) was proportional to the sigmoid steepness and inversely related to choice variability, *ρ* was the quantity ratio *q*_*B*_/*q*_*A*_ that made the animal indifferent between the two juices, and *ε* was a bias favoring one side over the other (specifically *ε* > 0 corresponded to a bias favoring the right side). Preliminary analyses indicated that the side bias was generally modest (|*ε*|<*ρ* in most sessions), but varied from session to session. Thus the analyses conducted here were restricted to sessions with a side bias |*ε*| ≤ 1 (see Livi et al. (2025)).We also fitted a reduced model excluding the side bias to compute *η* and *ρ*:

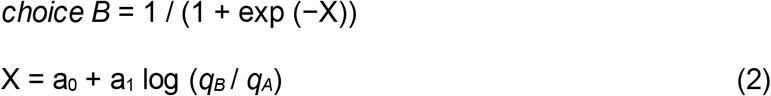

Again, from fitted parameters a_0_ and a_1_, we computed the choice accuracy *η* = a_1_ and the relative value *ρ* = exp(−a_0_/a_1_). Candidate variables possibly encoded by individual cells were defined based on the relative value measured in the same session.

### Analysis of neuronal responses

ΔF/F signals were examined in five time windows: *pre-offer* (600 ms preceding odor onset), *post-offer* (400-1000 ms from odor onset), *late delay* (1000-1600 ms from odor onset), *pre-choice* (600 ms preceding juice delivery onset), and *post-choice* (600 ms following juice delivery onset). An “offer type” was defined by two offered quantities. A “trial type” was defined by two offered quantities, their spatial configuration, and a choice. A “neuronal response” was defined as the activity of one neuron in one time window as a function of the trial type. Sessions typically included 10-12 offer types and 20-30 trial types. Neuronal responses were constructed by averaging the ΔF/F across trials for each trial type. Error trials in forced choices were excluded from the analysis. We also excluded from the analysis trial types with ≤2 trials.

As in previous studies (Kuwabara et al., 2020; Zhang et al., 2024), we assessed task-related modulation using a one-way ANOVA (factor: trial type) performed independently in each time window. A neuron was considered task-related if p<0.01 in at least one window. All subsequent analyses include only these neurons.

### LASSO regression

To determine a minimal set of decision variables necessary to explain each neuronal response, we employed LASSO regression. This analysis generalizes the approach used in our previous work, where each significant response was assigned to a single variable (Livi et al., 2025). Here, LASSO allowed us to obtain a sparse set of β coefficients for each neuron and each time window. For each session, we defined the same set of variables used in our previous studies, namely *position of A* (left or right; a variable that described the spatial configuration of the offers), *offer value A, offer value B, offer value ipsi* (the value of the offer presented on the ipsilateral side to the imaged neurons), *offer value contra* (the value of the offer presented on the contralateral side), *chosen value*, and *chosen side* (left or right) (Zhang et al., 2024; Livi et al., 2025).

Identifying what variable(s) is (are) encoded by each neuron is a noisy a process. To increase our accuracy in the identification process, we increased the signal-to-noise ratio by increasing the datapoints. Specifically, most FOVs (59 out 78) were imaged twice and neurons matched (see Zhang et al. (2024)). For successfully matched neurons, we used the activity of both sessions (Livi et al., 2025) to determine the coefficients. For each neuron, and each time window that passed the ANOVA, trial-by-trial ΔF/F values were regressed against the seven variables using LASSO, as implemented in Matlab 2021b by the `lasso.m` function. The model estimated the vector of β by minimizing the following objective function:

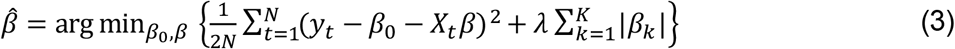

with *y*_*t*_ being the ΔF/F response on trial *t*, *X*_*t*_ the vector of predictor values for trial *t, N* the number of trials, *K* the number of variables, and *λ* is the regularization parameter controlling sparsity. Last, *β*_0_ was the intercept term, but because predictors were standardized to have mean zero and variance 1 within session, the intercept β0 was estimated but approached zero and is not interpreted.

LASSO regression (**Eq.3**) has the regularization parameter *λ*, which controls the penalty. We utilized the standard `lasso.m` implementation which evaluates a geometric sequence of *λ*. In our analysis, *λ* was selected independently for each neuron using a 10-fold cross-validation (**Fig.2A**). To favor model parsimony, and as per standard practice, we used the 1 standard error (1SE) rule (Hastie et al., 2009; Chen and Yang, 2021; Lee et al., 2022; Wang et al., 2024). In essence, we identified the *λ* that minimized the cross-validated means square error (MSE), and selected the largest *λ* within 1SE.

**Figure 2.**
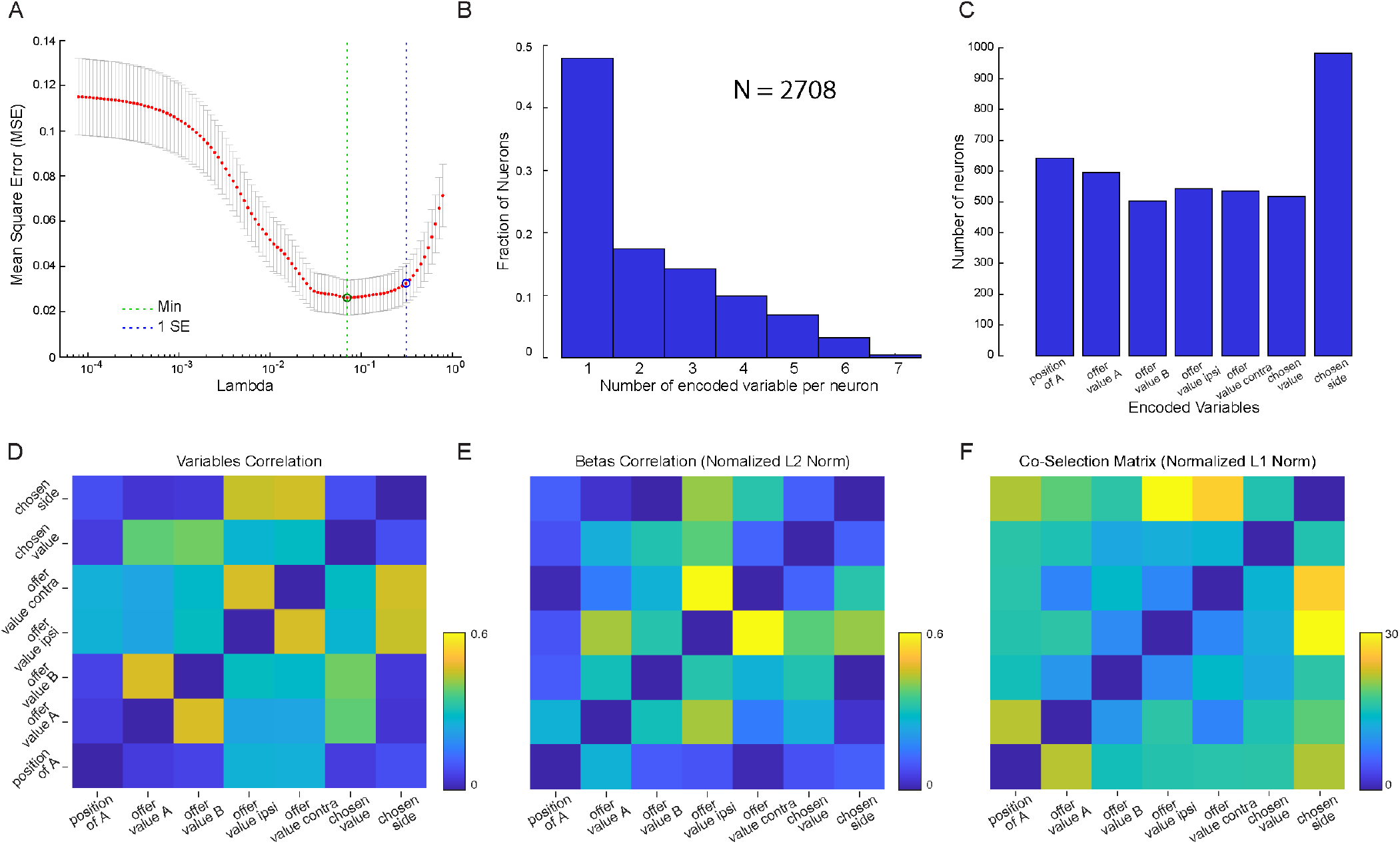
LASSO-based variable selection and task-variable correlation structure. **A**. λ selection applied to single-neuron activity. Mean squared error (MSE; y-axis) as a function of the regularization parameter λ. The red dots show mean ± SE across folds, for each value of λ. Vertical green dotted line: λ minimizing MSE. Blue dotted line: λ within one standard error (1SE rule). **B**. Distribution of encoded variables. Using LASSO, number of variables per neuron (N = 2708 task-related neurons; ≥1 ANOVA-significant window, p<0.01). Percentages: 48% encoded 1 variable, 17% encoded 2, 14% encoded 3, 10% encoded 4, 7% encoded 5, and 4% encoded ≥6. **C**. Frequency of variable encoding. Number of neurons representing each of the seven task variables under the 1SE λ solution. **D**. Correlation among task variables. Session-averaged Pearson correlation matrix across seven behavioral variables. Correlation was computed using task-variables absolute values. **E**. Correlation among LASSO β coefficients. Pearson correlation matrix of the magnitude (absolute value) of L2-normalized β coefficients (computed independently per time window). **F**. Co-selection matrix. β coefficients were binarized, then L1-normalized and used to quantify co-selection across variables. Each entry represents the number of times variable *i* and *j* were co-selected. The diagonal (self-selection) was set to 0 for visualization. Color scale reflects the number of occurrences (0–30).

**Figure 3.**
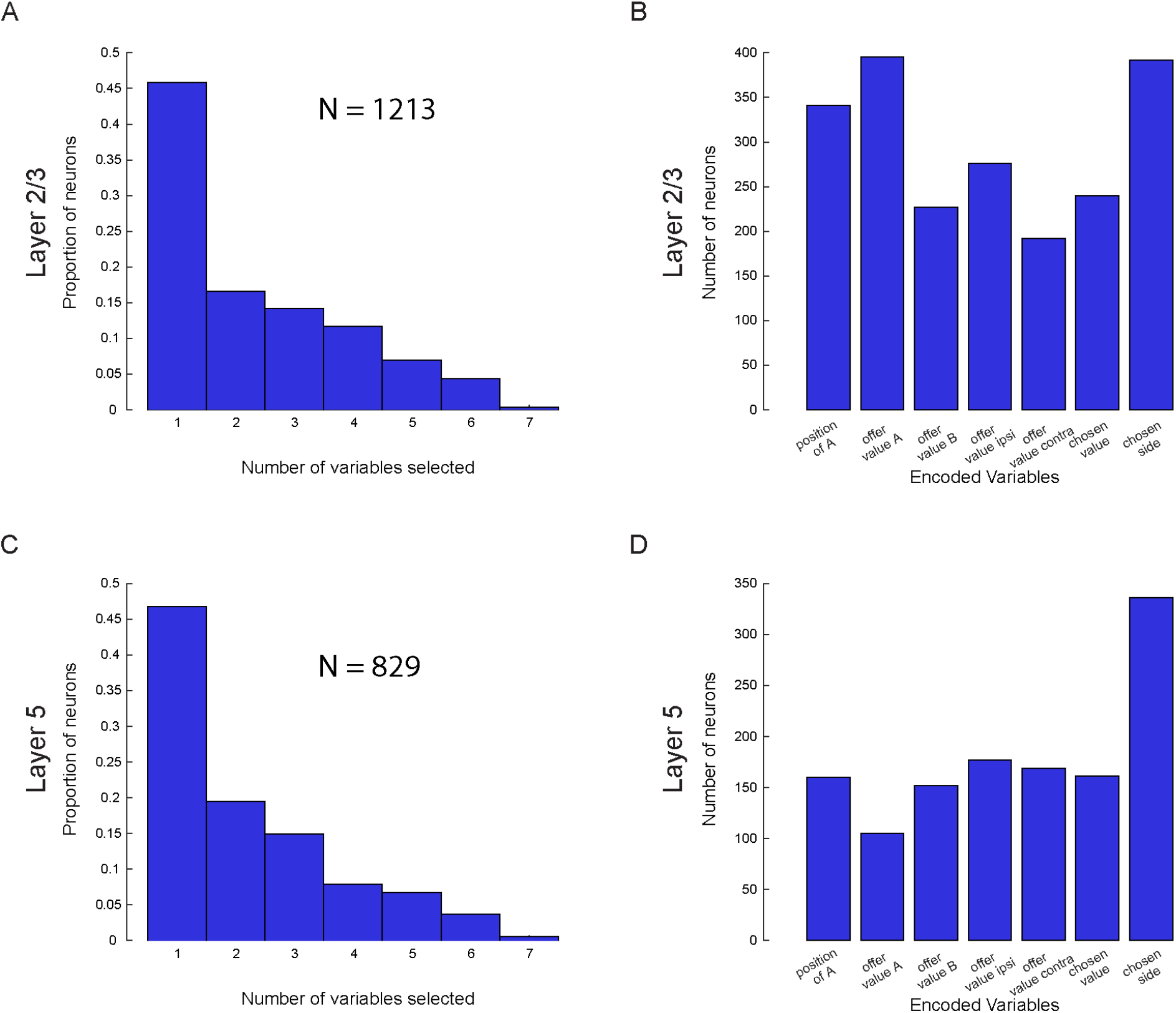
LASSO-based variable selection in layer 2/3 and layer 5 neurons. **A** Distribution of the number of task variables represented per neuron in layer 2/3, estimated using LASSO regression (N = 1213 task-related neurons; at least one ANOVA-significant time window, *p* < 0.01). Percentages of neurons representing 1 – 7 variables are: 46% (1), 17% (2), 14% (3), 12% (4), 7% (5), and 4% (≥6). **B** Frequency of variable representation in layer 2/3. Bars indicate the number of neurons with non-zero coefficients for each of the seven task variables under the 1SE λ solution. **C** As in **A**, but for layer 5 neurons (N = 829 task-related neurons). Percentages are: 47% (1), 20% (2), 14% (3), 8% (4), 7% (5), and 4% (≥6). **D** As in **B**, but for layer 5 neurons.

From these fits, a variable was considered “encoded” if its β coefficient was nonzero. In single-cell analysis, the number of encoded variables per neuron was defined as the count of all non zeros β coefficients over any time window passing the ANOVA. The correlation between the values of βs was computed using Pearson’s correlation as implemented in Matlab 2021b by the function `corr.m`, and computed for all task-related time windows. However, before computing the correlation, each vector of β coefficients was normalized with an L2 normalization. The beta vector for neuron *n* in time window *w* was:

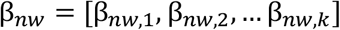

We then computed the L2 norm:

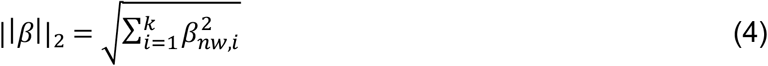

and we normalized each coefficient as follows:

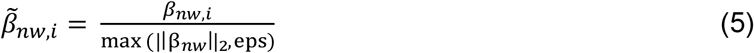

In **Eq.5**, *eps* is the small constant “eps” in Matlab that was used to avoid dividing by zero, *w* the time window and all other definition as in **Eq.3**. Correlation was computed on the absolute values of normalized β.

The co-selection matrix (**Fig.2F**) was computed as follows. For each neuron *n*, we identified all non-zero β, summed across time-windows and identified the logical vector of encoded variables:

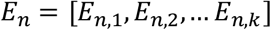

with *E*_*n*,*i*_ = 1 if variable *i* was encoded in ≥1 time window, and 0 otherwise.

To ensure comparability across neurons with different numbers of non-zero β, we normalized this vector using L1 normalization. L1 norm was computed as follow:

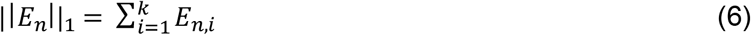

and the vector was normalized with:

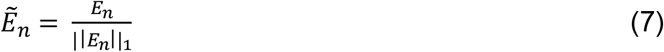

### Analysis of correlation between variables

Because several task variables in our design were not statistically independent, it was necessary to quantify the intrinsic correlations among them. If two variables are correlated at the level of trial structure, neurons may exhibit correlated LASSO coefficients simply as a consequence of this redundancy, rather than because they truly encode multiple variables. To correctly interpret the correlation among β weights, we therefore first computed the correlation structure of the task variables themselves (**Fig.2D**).

For each behavioral session, we extracted the trial-type vectors for each variable *X*_*k*_(e.g., offered values, chosen value, choice side). For every pair of variables (*k, j*), we computed the Pearson correlation coefficient:

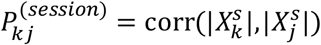

We then averaged these coefficients across all sessions:

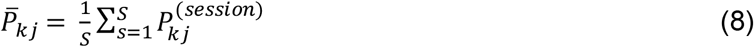

with S being the number of sessions. The resulting symmetric 7×7 matrix 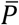 quantifies the typical correlation between each pair of task variables. This matrix provided the baseline level of expected correlation due solely to task structure. Consequently, any correlation observed among neural coefficients must be interpreted relative to this baseline.

### Decoding layer-specific population features using support vector machine (SVM)

We quantified layer-specific representation of task variables using support vector machine (SVM) classifiers (**Fig.4**). The analysis included only task-related neurons (≥1 time window passed the ANOVA threshold). For each neuron, we computed unregularized regression coefficients (“beta coefficients”) for four decision variables: *position of A, offer value A, offer value B*, and *chosen side*. We focused on this reduced set of four variables because it represents the minimal set of independent variables derived from the seven predictors used in previous analyses. Beta coefficients were extracted from three time windows –namely, late delay, pre-choice, and post-choice. We restricted the analysis to these three windows for two reasons. First, because we only regressed the neural activity in those time windows that passed the ANOVA, the remaining were set to zero and adding a fourth window would have increased the zeros excessively. Second, these three windows capture distinct computational stages of the decision process: offer evaluation (late delay), choice formation (pre-choice), and outcome processing (post-choice). Each neuron was therefore represented 0-as a 12-dimensional vector of β coefficients (4 variables × 3 time windows, concatenated). Before classification, we took the absolute values of β coefficients and non-finite entries (i.e., entries for which the ANOVA did not reach significance) were imputed with zeros, and each neuronal vector 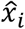 was self-normalized:

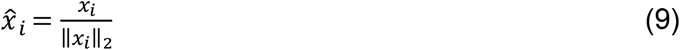

**Figure 4.**
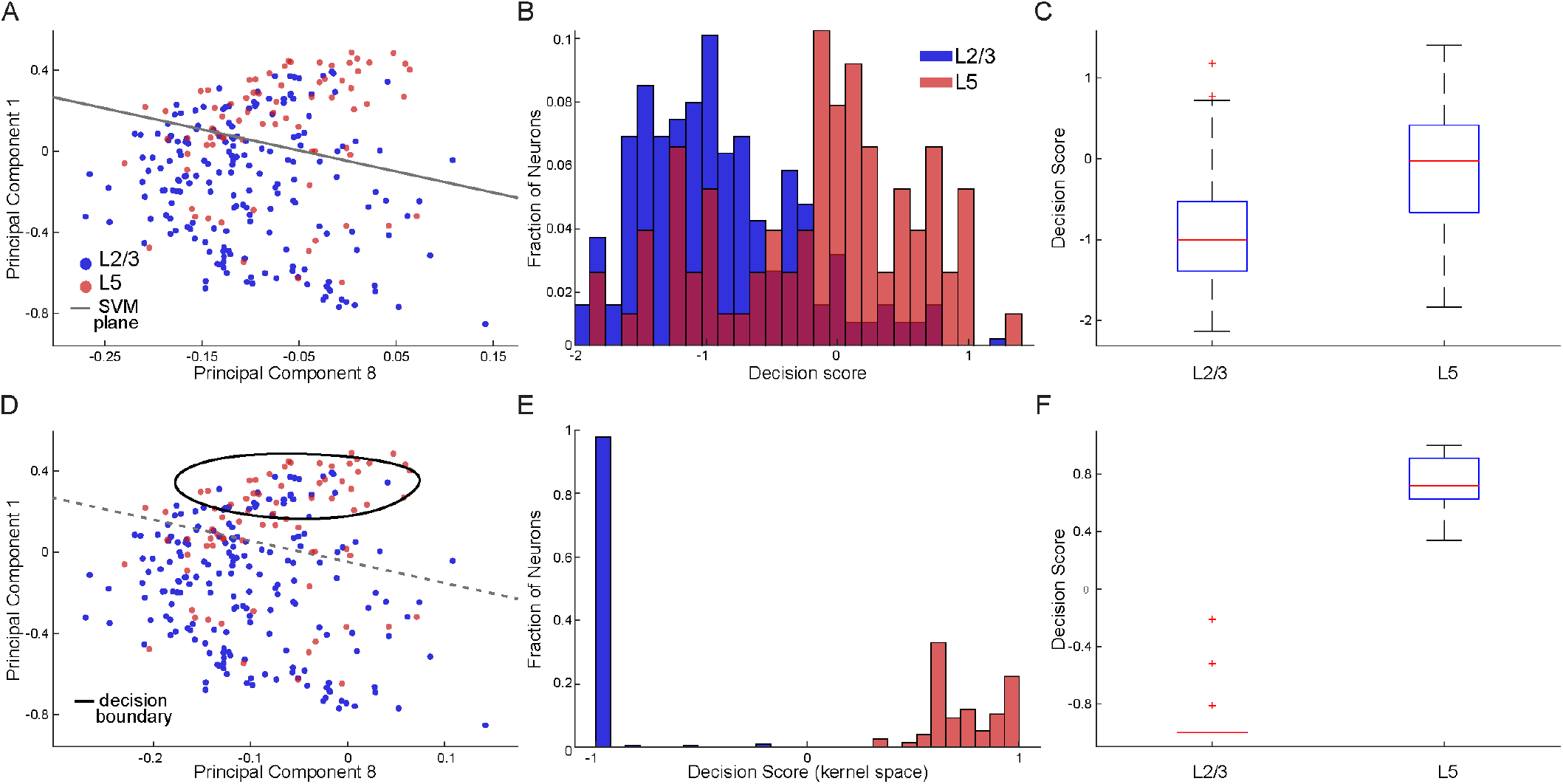
SVM classification of L2/3 and L5 populations from unregularized beta coefficients. **A**. Linear classifier. The two-dimensional projection of neuronal population unregularized beta coefficient matrix from L2/3 (blue) and L5 (red) cells is illustrated in PCA space. The x-axis corresponds to the principal component most aligned with the linear SVM normal vector (here PC 8, explained 1.8% of the variances), while the y-axis shows PC1 (explained 35.4% of the variances). Each point represents a single neuron’s beta coefficient vector after taking absolute values and within-neuron normalization across task variables and time windows. The solid black line marks the linear decision boundary (score = 0) computed from the cross-validated linear SVM. Model performance (mean ± SD across 5-fold CV): ACC = 0.78 ± 0.03, AUC = 0.75 ± 0.04, *p* < 0.001 for both ACC and AUC vs. label-shuffled null (1,000 permutations). **B**. Linear decision scores. The two histograms illustrate the linear SVM decision scores (projection onto the SVM weight vector *w*) for neuronal populations in L2/3 (blue) and L5 (red). Positive scores indicate stronger L5-like encoding; negative scores indicate L2/3-like patterns. The partial overlap between distributions reflects neurons with intermediate mixed-layer representations. **C**. Boxplots summarizing linear decision scores for the two populations. The center red line indicates the median; boxes mark the interquartile range (25th – 75th percentile); whiskers extend to 1.5× the interquartile range; and individual points denote outlier neurons with unusually high or low decision scores. The overall distribution confirms that L5 neurons tend to have higher (more positive) linear SVM scores than L2/3 neurons. **D**. Nonlinear classifier. Data are shown in the same PCA projection as in **A**, but with the nonlinear (RBF) SVM boundary superimposed. The black contour outlines the region where the RBF decision function equals zero (score = 0), corresponding to the nonlinear classification boundary in the original feature space, while the dashed black line shows the linear boundary generated from panel **A** for reference. The RBF model achieves ACC = 0.74 ± 0.02, AUC = 0.81 ± 0.04, *p* < 0.002 for ACC and *p* <0.001 for AUC (1,000 permutations), indicating improved separability and capturing curved manifolds separating L2/3 and L5 activity patterns. **E**. Nonlinear decision scores. Unlike the linear projection, RBF scores saturate near −1 and +1, reflecting confident classification of most neurons. **F**. Boxplots of RBF SVM classifier decision scores for L2/3 and L5 neurons. Compared to the linear model, the RBF SVM classifier yields greater separation and smaller overlap between layer distributions, consistent with improved discrimination in a nonlinear feature space.

This normalization ensures that the classifier emphasizes directional encoding patterns rather than absolute response magnitude, facilitating comparison across heterogeneous firing rates and signal amplitudes. We trained binary SVM classifiers to discriminate between neurons recorded in L2/3 and L5. For the linear SVM, the decision rule was defined in terms of

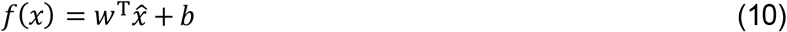

where *w* is the weight vector normal to the separating hyperplane and *b* is the bias term. Predicted labels were obtained as 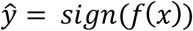, and the decision score used for plotting and Receiver Operating Characteristic analysis (ROC) corresponds to the signed distance from the hyperplane. The model was optimized by minimizing the standard hinge-loss objective with soft-margin penalty C:

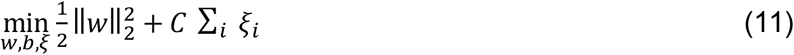

with *ξ*_*i*_ the slack term of the SVM subject to:

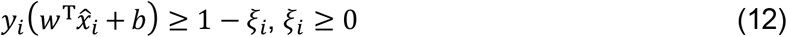

where *y*_*i*_ = 0 for L2/3 and *y*_*i*_ = 1 for L5 cells. To account for class imbalance, we applied asymmetric misclassification penalties, scaling the L5 error cost by a factor equal to 1.24. This value was chosen since returned the highest SVM classifier performance. This weighting ensured balanced contribution of both populations to the decision boundary.

For the nonlinear SVM, we used the radial basis function (RBF) kernel:

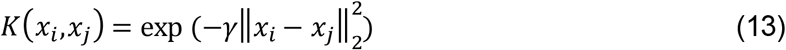

with 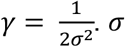 determines how quickly *K* decays with distance between two neuronal feature vectors *x*_*i*_ and *x*_*j*_. Use of a kernel implicitly projects data into a higher-dimensional feature space where nonlinear boundaries become linearly separable.

The resulting nonlinear decision function was:

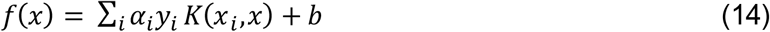

The multipliers *α*_*i*_, (*α*_*i*_ > 0) determine how much each neuron contributes as a support vector to the final decision boundary, while *C* sets the maximum allowed influence of any single sample, thereby controlling the model’s capacity and tolerance to margin violations. The RBF classifier captures curved separations in the representational geometry that are not well described by a single linear projection.

Model performance was evaluated using 5-fold cross-validation: the dataset was partitioned into five non-overlapping folds, with four folds used for training and one for testing in each iteration. Mean classification accuracy (ACC) and area under the ROC curve (AUC) were computed across folds.

Statistical significance was assessed via 1,000-fold label-shuffled permutation tests, in which the neuron-layer labels were randomized while maintaining the same 5-fold cross-validation structure. This generated a null distribution of ACC and AUC values against which the observed performance was compared.

To visualize the decision geometry, neuronal feature vectors were projected onto the first principal component aligned and the one most aligned with the linear SVM normal vector, forming a 2-D subspace that best captures layer separation. Decision boundaries for both linear and RBF models were plotted as the locus where the classifier score equals zero (*f*(*x*) = 0).

### Activity profiles and strength of the encoding

The general construction of this analysis follows the same logic previously described (Livi et al., 2025). The only difference was that here each neuron contributed to every variable with a continuous weight derived from LASSO coefficients, rather than contributing to a single variable. The dataset included all neurons classified as task-related (≥1 time window passed the ANOVA threshold), and had at ≥1 non-zero β. Instead of assigning each neuron to a single variable category, we used the L1-normalized selection vector 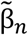. In essence, for each neuron we first summed across time windows, computed the ||β_*n*_||_1_ as follow:

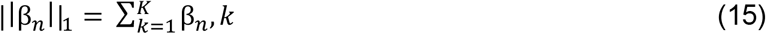

then, we normalized 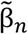:

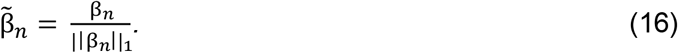

Importantly, we preserved the signs since, in this analysis, neurons with positive and negative encoding were treated differently (see below). Specifically, first cells with positive and negative encoding were kept separate (**Fig.5** for L2/3, **Fig.6** for L5), and then pooled together (**Fig.7**) by reversing the order of the quantiles for the negative sign (i.e., quantile 1 became quantile 3, and quantile 3 became quantile 1). The activity was aligned to both the offer onset and the choice. For this analysis, the neural contribution to *offer value A* and *offer value B (offer value A*|*B)*, and *offer value I* and *offer value C (offer value I*|*C)* were pooled (as in previous work). Cells from L2/3 and L5 were examined separately. For binary variables (*position of A* and *chosen side)*, trials were divided in *ipsilateral* and *contralateral*, and the side eliciting the highest response defined as *encoded*, the other as *not encoded*. For continuous variables (*offer value A*|*B, offer value I*|*C, chosen value*), trials were divided in three sets such that the encoded variable would be low, medium, or high (three quantiles of equal width).

**Figure 5.**
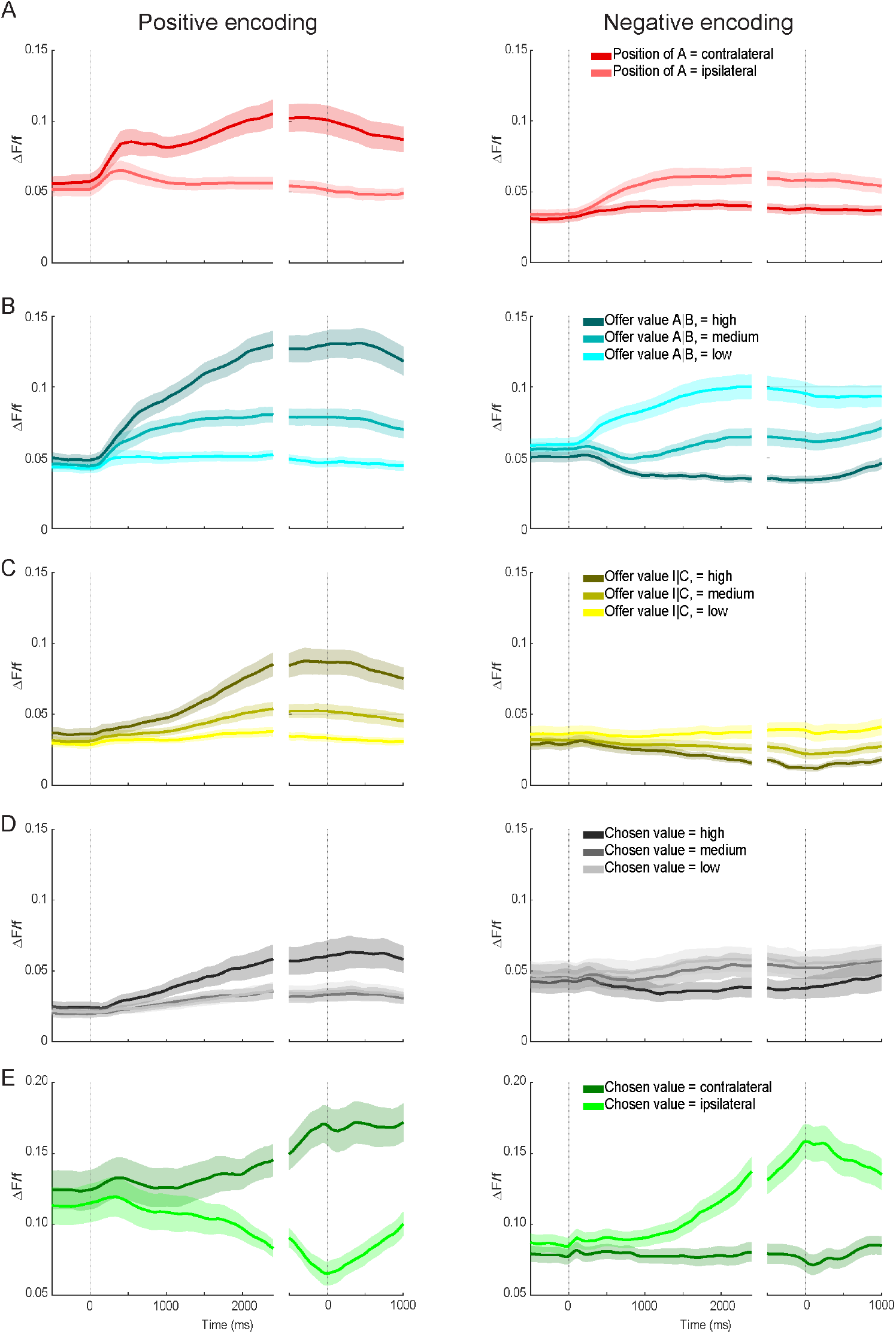
Activity profiles of layer 2/3 neurons. **A** Cells encoding the variable *position of A*. Neurons were separated based on the sign of encoding: cells showing higher activity when position of A was contralateral (“positive encoding”, left panels) and cells showing higher activity when position of A was ipsilateral (“negative encoding”, right panels). For each neuron, trials were grouped according to whether juice A was offered on the ipsilateral or contralateral side. Trials were aligned separately to offer onset and juice delivery (two vertical dashed lines). Neural activity (ΔF/F) was computed for each neuron and averaged across neurons to obtain population activity profiles (see **Methods**). Lines indicate the mean; shaded areas represent running ±2 s.e.m. **B** Cells encoding the variable *offer value A*|*B*. Neurons encoding *offer value A* and *offer value B* were pooled and separated by encoding sign (positive, left panels; negative, right panels). For each neuron, trials were divided into low, medium, and high value bins (see **Methods**), and activity profiles were computed and averaged across neurons for each bin. **C** Cells encoding the variable *offer value I*|*C*. **D** Cells encoding the variable *chosen value*. **E** Cells encoding the variable *chosen side*. All times are in milliseconds. All other conventions are as in panel **A**.

**Figure 6.**
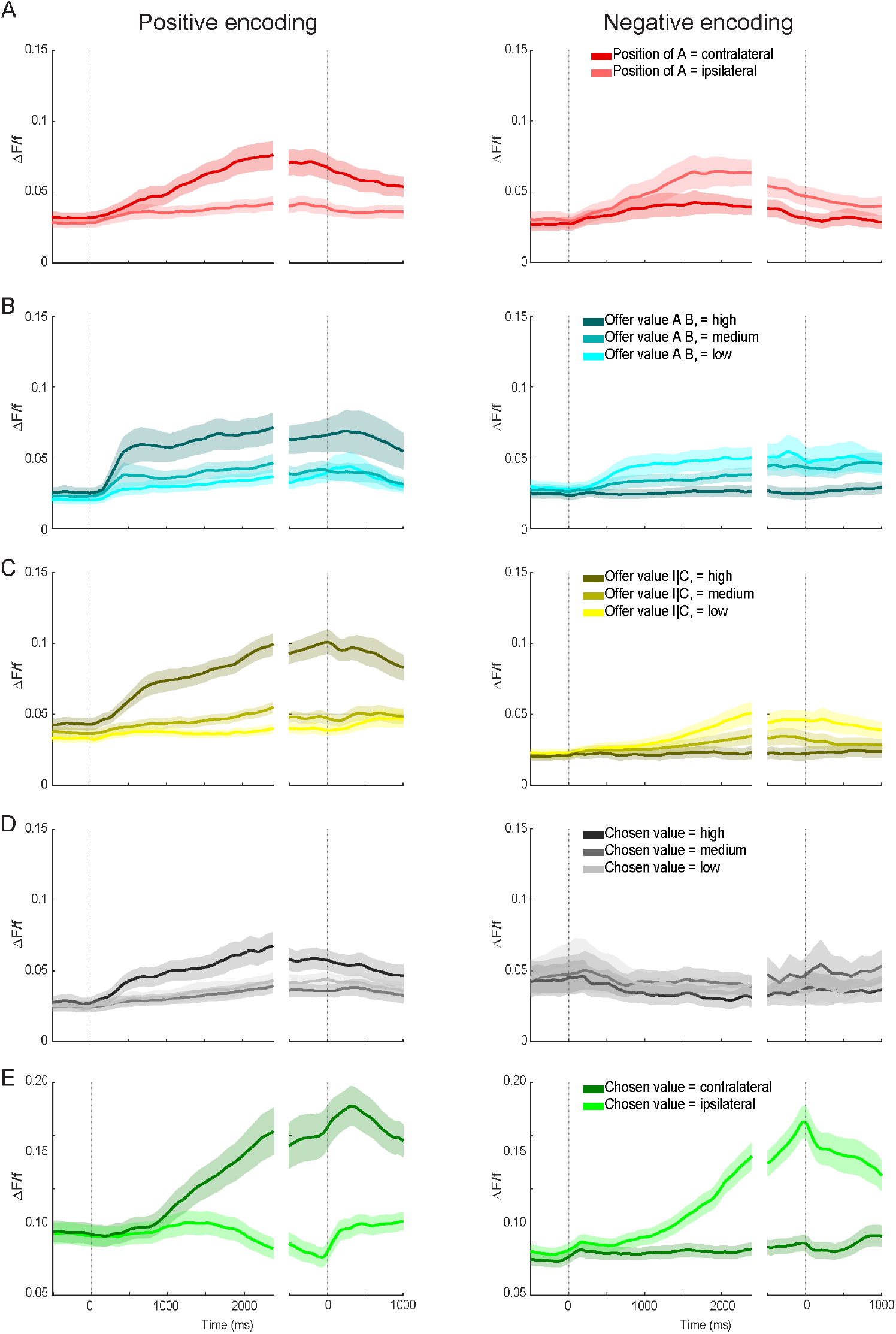
Activity profiles for layer 5. All conventions as in **Figure 5**.

**Figure 7.**
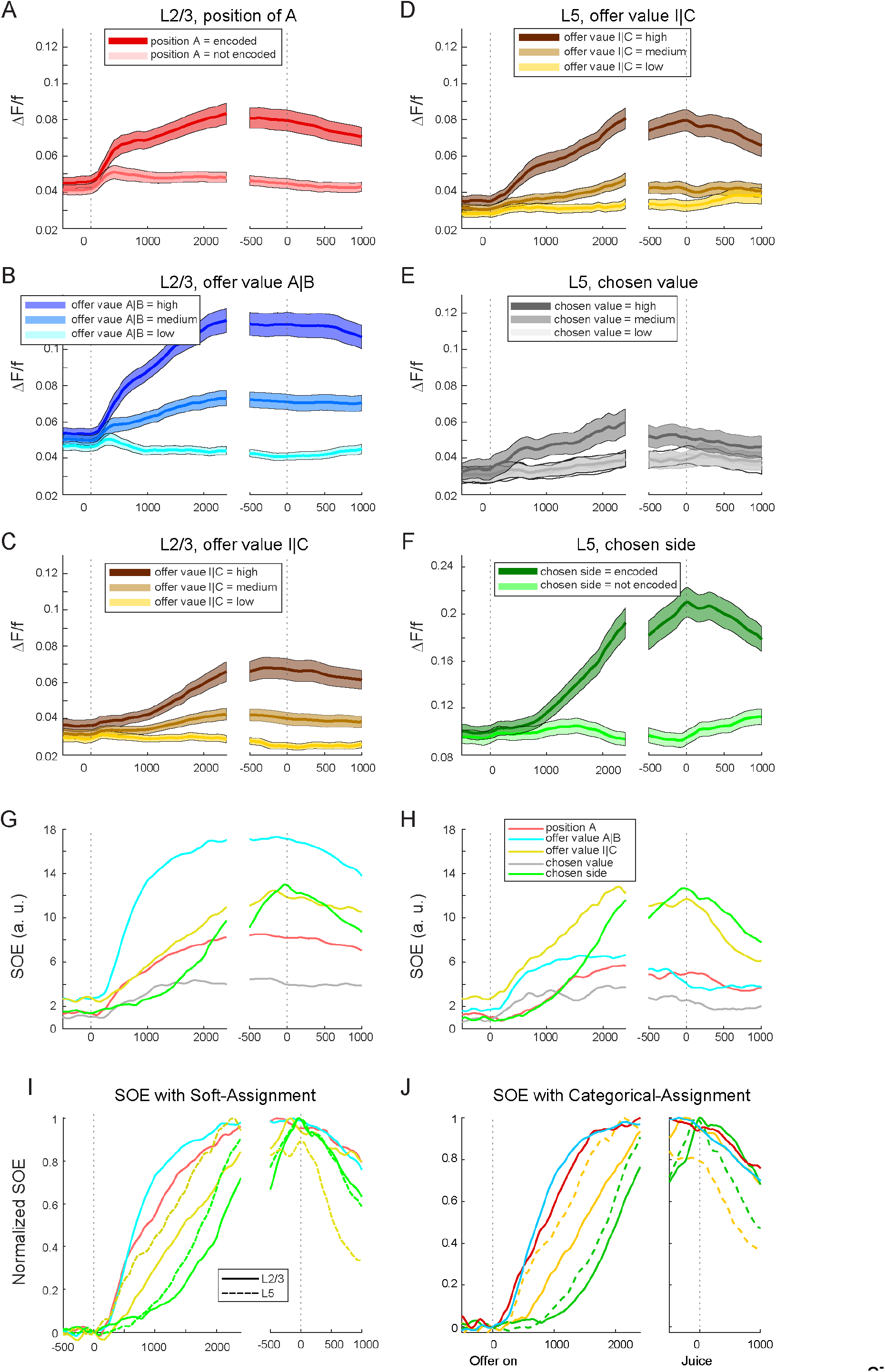
Population activity profiles and strength of encoding (SOE). **A**. Position of A (L2/3). For each neuron, trials were split according to whether juice A appeared to the preferred side (highest response) or non-preferred side (lowest response). Activity traces (ΔF/F) were aligned to offer onset and juice delivery (two vertical dashed lines), weighted by each neuron’s L1-normalized β coefficient, and averaged across neurons (N = 2708). **B**. Offer value A|B (L2/3). Neurons contributing to offer value A or B were pooled. Trials were sorted into low/medium/high quantiles of the relevant offer value; for neurons with a negative sign the quantiles were flipped to be combined. Weighted activity profiles were computed and averaged. **C**. Offer value I|C (L2/3). **D**. Offer value I|C (L5). **E**. Chosen value (L5). **F**. Chosen side (L5). Panels **A – F** Mean ± s.e.m. is plotted. **G – H**. Strength of encoding (SOE). SOE traces for L2/3 (G) and L5 (H), computed from absolute β coefficients normalized by the L1 norm across variables at each time bin. Positive and negative encodings were combined. **I**. Normalized SOE. Each SOE trace was baseline-subtracted (at offer onset) and normalized by its peak amplitude. Continuous lines = L2/3; dashed lines = L5. As in categorical analyses, offer value A|B and position of A emerged earliest in L2/3, offer value I|C appeared earlier in L5, and chosen side rose latest – mirroring temporal hierarchies from our previous categorical work. In all plots, first vertical dashed line is the offer on, second is animal’s choice. **J** As in **I** but the analysis was done with a categorical assignment in our previous work. A different color tone is used to indicate that this panel was generated using categorical assignment.

For each cell and each set of trials, we averaged ΔF/F across trials. For each variable, layer, and sign of encoding, we then computed the weighted population activity profile:

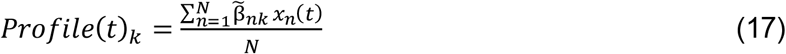

where *x*_*n*_(*t*)is the mean ΔF/F trace. Then we then computed the mean, standard deviation, and standard error across the population, for each value of the encoded variable.

Based on the activity profiles, we also computed the strength of the encoding (SOE) (**Fig.7G, H**). We proceeded exactly as in our previous work (Livi et al., 2025). Specifically, SOE was computed for each variable and for each sign of encoding, and was a function of time (*t*) in the trial. First, we computed the activity range (AR). For binary variables:

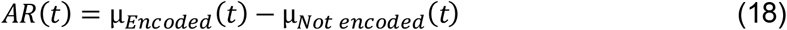

while for trinary variables, we regressed the weighted mean responses of three quantiles to the specific variables and defined:

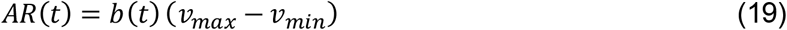

with *b(t)* the regression slope at time *t* and *v* the x position during the regression (i.e., the first quantile and third). Then, SOE was defined as:

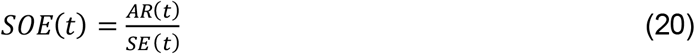

and SE defined as the Standard Error. We combined positive and negative encoding populations with:

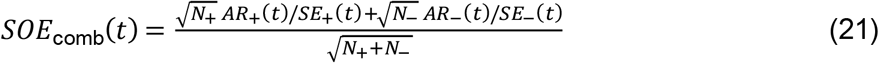

where +/-refer to populations with positive and negative encoding, respectively. Last, for temporal comparison, we normalized the SOE by subtracting the baseline, defined as the activity at the beginning of the offer on, and divided by its peak:

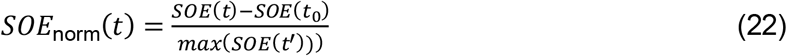

*t’ is* the max value of SOE in the entire analysis window (**Fig.7I**).

### Granger Causality Analysis

GCA followed the Multi Variate Granger Causality toolbox, implemented as previously described (Livi et al., 2025). ΔF/F time series were concatenated across trials and modeled with VAR processes.

Model order was determined using Akaike Information Criteria (AIC) (lags 1–5 across sessions).

Pairwise conditional GCA was computed, and significant effects (p < 0.01, false discovery rate corrected) yielded a binarized directed adjacency matrix per session.

#### Connections within and between cell groups with soft assignments

To analyze connectivity within and between functional groups, we adapted our previous categorical (hard-assignment) approach to a soft-assignment framework based on LASSO coefficients. Instead of assigning each neuron to a single variable category, we used the L1-normalized selection vector 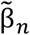 as we did for the analysis of profiles (see **Eq.15, 16**), but in this case after taking the absolute value of the β coefficients.

Normalization ensured that the weights summed to 1 ensuring that each neuron contributed equally. For each session (example in **Fig.8A**), GCA adjacency matrix was as in our previous work (Livi et al., 2025) and then combined with the soft assignment. *Connections Count* (**Fig.8B**) for any pair of variables (*k*_*i*_,*k*_*j*_):

**Figure 8.**
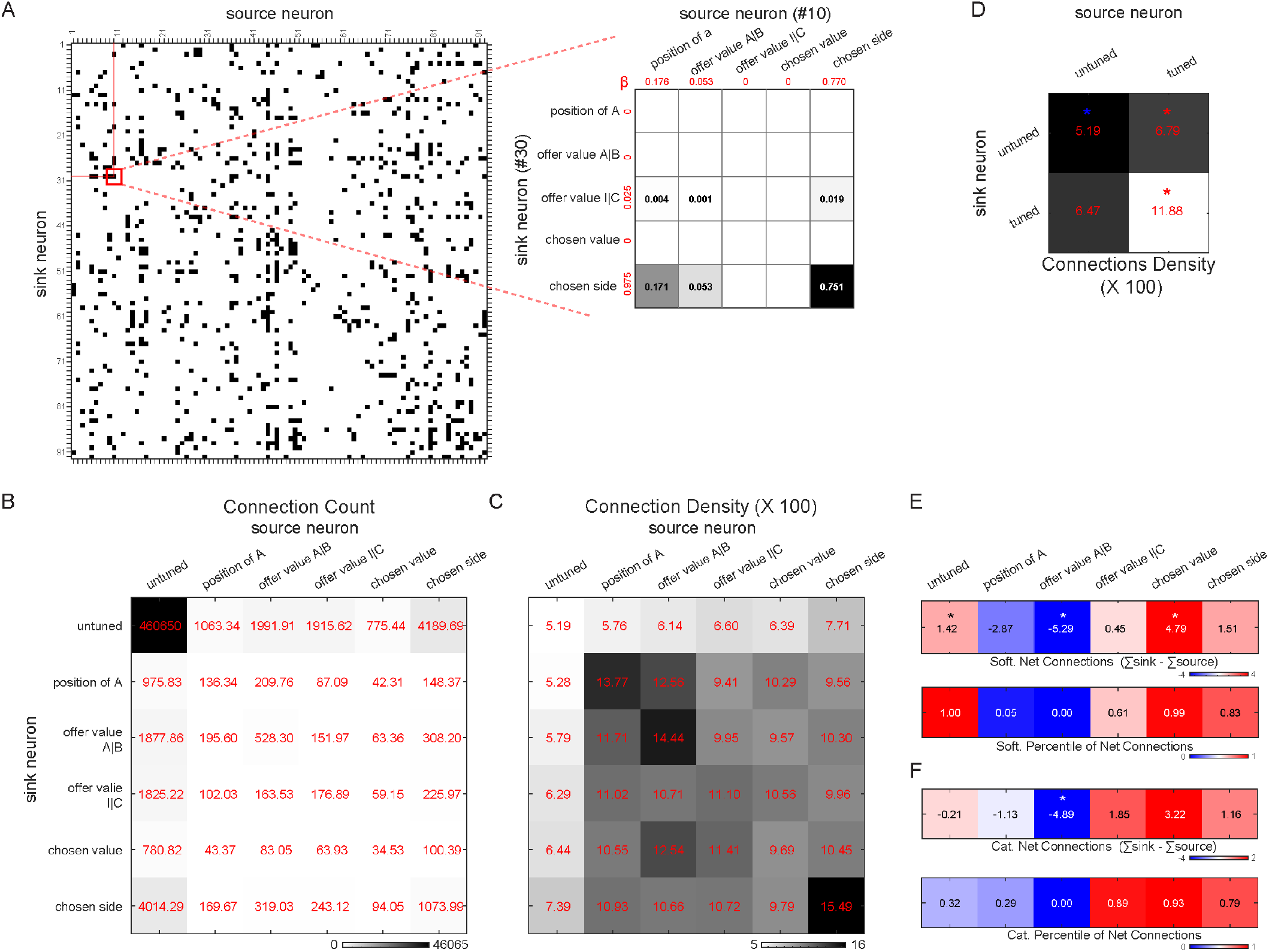
Granger-causality with soft assignment. **A** Example session. Left: binarized Granger-causality adjacency matrix (columns = source neurons; rows = sink neurons). Black squares indicate significant directed influences (*p* < 0.01, corrected). Right: zoomed-in example of a single significant connection (red box on the left), illustrating how one neuron-to-neuron GC connection is redistributed into variable-to-variable contributions according to the L1-normalized β-vectors (numbers in red) of the source and sink neurons. **B** Connection counts. Total number of GC connections across the population for each ordered pair of variables, obtained using soft assignment. **C** Connection density. Each entry in **B** is divided by the total number of possible connections (values ×100 for visualization). **D** Connection density for tuned versus untuned neurons. GC connections were marginalized into a 2 × 2 matrix based on whether source and sink neurons were tuned (i.e., encoded at least one task variable) or untuned. Each entry represents the density of directed connections (×100) between the corresponding source – sink categories. Asterisks indicate entries that significantly deviate from the bootstrap null distribution (N = 10,000; *p* < 0.05, corrected). **E** Soft-assignment net connections and bootstrap percentiles. Top: net GC connections (row sum minus column sum) computed from soft assignment. Bottom: percentile ranks of net connections relative to the bootstrap null distribution (N = 10,000). **F** Categorical net connections and bootstrap percentiles. Same analysis as in **E**, but using categorical GC assignments from Livi et al. (2025).

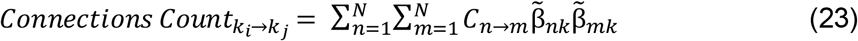

with *C*_*n*→*m*_ being the binarized GCA adjacency matrix for neuron *n* and *m*, 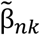 and 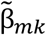 the normalized betas for neuron *n* and *m*, and variable *i* and *j*. Next, we defined the *Possible Connections* for any pair of variables (*k*_*i*_,*k*_*j*_):

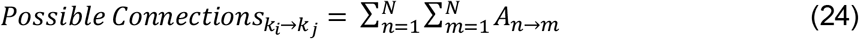

with *A*_*n*→*m*_ being the adjacency matrix as if all connections were 1 but *n* ≠ *m*. To compute population data, all the *Connections Count*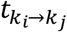 across all sessions were summed, similarly also *Possible Conenctions*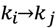 across sessions were summed, and density (**Fig.8C**) computed as follows:

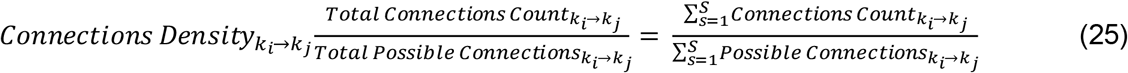

Then, we computed the *Reduced Connection Density* (**Fig.8D**) reducing full *Connection Density* to a 2×2 table (*untuned, tuned*). Using a similar approach, we also computed the *Net Connection Density* (**Fig.8E**). In this case, for each variable we computed the sum of all *Sink Connection Density* (row) and subtracted that value by the sum of all *Source Connection Density* (column). To assess the significance of *Net Connection Density*, we performed a bootstrap analysis (p<0.05, uncorrected, two tailed). Essentially, we shuffled the position of the significant connections in the adjacency matrix, while preserving the total sum of connections. We repeated this procedure N = 10,000 times, computed the distribution of shuffled *Connections Count*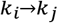, computed the *Net Connection Density* and compared the actual data to the shuffled distribution.

## Results

### Choice task and 2P calcium imaging of OFC

Mice were head-restrained and positioned under a two-photon (2P) microscope (**Fig.1A**). Animals were trained to perform a juice choice task analogous to those used in primate studies (Padoa-Schioppa and Assad, 2006). In each session, animals chose between two juice flavors, labeled A and B (with A preferred to B). The experimental design was adapted to mice in the sense that offers were represented by olfactory stimuli instead of visual cues (Kuwabara et al., 2020; Zhang et al., 2024; Livi et al., 2025). Olfaction is an ethologically strong sensory modality for mice, and odors reliably convey abstract information such as reward identity and quantity (Millman and Murthy, 2020). In each trial (**Fig.1B**), two odors were delivered from the left and right ports; the odor identity indicated the juice type, and odor concentration indicated the juice quantity. Across trials, we varied the offered quantities of two juices (A preferred to B) and counterbalanced the spatial configuration of the offers.

Behavioral choices were fitted with a logistic regression (see **Methods**), yielding three parameters: relative value (ρ), choice accuracy (η), and side bias (ε). In essence, the relative value captured the quantity ratio that made the animal indifferent between the two juices (flex of the sigmoid**; Fig.1C**); the choice accuracy captured the degree of choice consistency (sigmoid steepness**; Fig.1C**); the side bias captured any systematic preference for one side versus the other. Sessions satisfying our behavioral criteria – i.e., high accuracy and small side bias; see **Methods** – were included for neural analyses.

We imaged calcium dynamics from neurons in the lateral orbital area (here referred to as OFC) of 14 animals while they performed the choice task. The cortical volume was sampled using multiple fields of view (FOVs) positioned along the z-axis to avoid duplicating neurons across planes. Neuronal signals extracted with CaImAn were expressed as ΔF/F. In total, we imaged 78 FOVs. Neurons matched across sessions (N = 2445) were combined, yielding a final dataset of 9347 unique neurons.

Neuronal activity was examined in five time windows. A trial type was defined by the combination of offered quantities, spatial configuration, and choice. A neuronal response was defined as the activity of one neuron in one time window, as a function of the trial type. Each neuronal response was screened with a one-way ANOVA (factor: trial type; p < 0.01; see **Methods**). Of all recorded neurons, N = 2708 were modulated by the trial type in at least one time window, and were thus identified as task-related. Each neuronal response passing the ANOVA criterion was regressed against seven task variables identified in previous studies and accounting for >85% of OFC responses, namely *position of A, offer value A, offer value B, offer value ipsi, offer value contra, chosen side*, and *chosen value* (Zhang et al., 2024; Livi et al., 2025).

### Neuronal encoding of decision variables in OFC with LASSO

Neurophysiology studies in primates and rodents have shown that neurons in OFC represent individual offer values, the binary choice outcome, and the chosen value. The fact that these variables capture both the input and the output of the decision process, together with a series of corroborating results, has suggested that OFC hosts a neural circuit in which economic decisions are formed.

In our previous study (Livi et al., 2025), each neuron was classified as encoding a single variable using the following procedure. First, each neuronal response passing the ANOVA criterion was regressed against each variable (unregularized regression), yielding an R^2^ value. When the regression slope was statistically indistinguishable from zero, or when the response did not pass the ANOVA, we arbitrarily set R^2^ = 0. Second, for each cell and each variable, we summed the signed R^2^ values across time windows (the sign was given by the sign of the regression slope) and computed the absolute value R^2^_total = |Σ_time windows (slope sign × R^2^)|. Third, we assigned the cell to the variable yielding the largest R^2^_total.

Using this approach, we examined the laminar organization of the decision circuit in OFC (Livi et al., 2025). Within this framework, we showed that decision variables are differentially represented across layers – juice-specific offer values and their spatial configuration are predominant in L2/3, whereas spatial offer values, chosen side, and chosen value are predominant in L5. Functional connectivity between cell groups, assessed through GCA, supported the notion that neurons encoding individual offer values provide the primary input to a decision circuit. Moreover, the temporal dynamics of neural signals in the two layers indicated a combination of feedforward and feedback processes and pointed to L5 as the locus of winner-take-all value comparison. Taken together, these results provided a more detailed understanding of the neural mechanisms underlying economic decisions. However, in light of the recent findings in non-human primates (Crimmins et al., 2025), one concern is whether these results would be stable even if neurons in OFC exhibit some degree of mixed selectivity.

To determine whether these results are affected by restricting each neuron to a single variable, we relaxed this condition by performing a LASSO analysis of the variable(s) encoded by individual responses. LASSO regression penalizes nonzero coefficients using an L1 penalty on β-coefficients weighted by a scalar denoted λ; this produces sparse solutions with zero, one, or more than one nonzero β-coefficients per neuron. For each task-related neuron (N = 2708), we: (i) collected all ANOVA-significant time windows (from one or two registered sessions); (ii) fit a single LASSO model (see **Methods**); (iii) performed 10-fold cross-validation; and (iv) selected λ using the 1-SE rule (**Fig.2A**). This procedure yielded seven β-coefficients per response, of which any number might be zero.

The number of variables encoded per neuron was defined as the number of nonzero β-coefficients (**Fig.2B**). The resulting distribution was: 48% encoded one variable, 17% two, 14% three, 10% four, 7% five, and 4% six or seven. This also revealed that chosen side was the most frequently encoded variable (**Fig.2C**), consistent with prior analyses based on single-variable assignments (Zhang et al., 2024; Livi et al., 2025).

At first glance, the distribution in **Fig.2B** suggests two populations: a categorical group (single-variable neurons) and a second group potentially exhibiting mixed selectivity (neurons encoding more than one variable). However, because the task variables were not independent, apparent multi-variable encoding might arise trivially from correlations intrinsic to the task structure. As a result, mixed selectivity must be evaluated relative to the intrinsic level of correlation among variables (Onken et al., 2019; Crimmins et al., 2025). To investigate whether this might be happening, we quantified correlations among the task variables themselves (**Fig.2D**) and compared them with correlations among LASSO β-coefficients after L2 normalization (**Fig.2E**). Inspection of these results indicates that neural correlations closely follow, but do not perfectly overlap with, correlations among task variables.

To gain further insight into whether variable selection is entirely driven by the task, we examined how variables tended to be co-selected. For each neuron, we created a logical vector of non-zero β-values, normalized it by its sum, and computed variable co-selection via the outer product. In the population-level co-selection matrix (**Fig.2F**), with the diagonal removed, the highest co-selection entries were observed between *offer value-ipsilateral* or -*contralateral* (first and second highest, respectively) with *chosen side*, suggesting entanglement between representations of *offer value ipsilateral/contralateral* and *chosen side*. This result is notable given that they do not simply follow the most frequently selected variables overall. In descending order, the most frequently selected variables were *chosen side* (N = 128.7), *position of A* (N = 106.4), *offer value ipsilateral* (N = 92.1), *offer value A* (N = 90.6), *offer value contralateral* (N = 86.9), *chosen value* (N = 84.7), and *offer value B* (N = 75.9).

In summary, applying LASSO regression provided (i) a substantive test of whether our previous results hinge on restricting neurons to single decision variables, and (ii) an estimate of the prevalence of multi-variable encoding. Approximately half of all neurons (48%) were best explained by a single variable, whereas the remaining neurons (52%) exhibited mixed selectivity. Similar proportions were observed when analyzing neurons separately in superficial (L2/3) and deep (L5) layers (**Fig. 3**; see below). Crucially, we caution against over-interpreting the distribution of non-zero regression coefficients. This distribution is shaped by task structure, correlations among predictors, neuronal noise, and other methodological factors. Accordingly, we defer broader conclusions about the nature of OFC representations to ongoing work (Crimmins et al., 2025). In the following sections, we assess whether previous conclusions regarding laminar architecture, functional organization, and circuit dynamics remain valid when single neurons are allowed to exhibit mixed selectivity.

### Layer 2/3 and layer 5 represent different mixtures of decision variables

Our dataset included neurons imaged from different cortical layers. To test whether the results described above were consistent across layers, we replicated the analyses of **Fig.2B,C** separately for L2/3 (**Fig.3A,B**) and L5 (**Fig.3C,D**). The distributions of the number of variables encoded by each neuron (**Fig.3A,C**) were remarkably similar across layers, suggesting that the type of encoding (i.e., categorical vs mixed selective) does not depend on cortical layer. By contrast, the specific variables encoded by neurons differed between L2/3 (**Fig.3B**) and L5 (**Fig.3D**). Consistent with our previous results where neurons were encoding at most one variable (Livi et al., 2025), neurons in L5 predominantly encoded *chosen side*, whereas neurons in L2/3 predominantly encoded *offer value AB*.

Using categorical assignments, we previously showed that decision variables are differentially represented in L2/3 and L5, based on odds ratio analyses (Livi et al., 2025). Here, we tested whether these layer-specific differences persisted when allowing neurons to encode linear combinations of variables. We addressed this question using support vector machine (SVM) decoding. In essence SVM decoding tests whether the organization of encoding vectors defined in a high-dimensional space differs systematically between layers. We therefore examined whether population activity patterns differ between L2/3 and L5 using SVM classifiers trained on task-related neurons. Analyses were limited to three time windows: late delay, pre-choice, and post-choice (see **Methods**).

For each neuron, we performed an unregularized multivariate linear regression across trials against four variables – position of A, offer value A, offer value B, and chosen side – yielding a 12-dimensional β-vector (4 variables × 3 time windows). A linear SVM trained on these vectors achieved robust above-chance discrimination between layers (5-fold cross-validated accuracy = 0.78 ± 0.03, AUC = 0.75 ± 0.04; p < 0.001 for ACC and AUC, 1,000 permutations), indicating that L2/3 and L5 neurons encode partially distinct task representations even without enforcing single-variable representations.

Projection of the neural population onto a two-dimensional PCA subspace, with one axis chosen to be that best aligned to the SVM normal vector, revealed a partial separation between L2/3 and L5 neurons along the linear decision axis. This separation on held-out data indicates that layer identity is embedded within shared dimensions of task-related variance (**Fig.4A-C**).

We also examined layer differences using a nonlinear SVM (RBF kernel; see **Methods**). Classification performance improved (accuracy = 0.74 ± 0.02, AUC = 0.81 ± 0.03; p < 0.002 for ACC and p < 0.001 for AUC, 1,000 permutations), and decision-score distributions became sharply bimodal (**Fig.4D-F**).

This suggests that curved decision boundaries more accurately capture the local structure of neuronal manifolds in high-dimensional space, and that the apparent overlap under linear decoding arises from curvature in the underlying neural manifold.

In summary, similarly to what we observed in our previous work under the constraint of categorical encoding (Livi et al., 2025) is relaxed, neuronal populations in L2/3 and L5 are found to represent different mixtures of decision variables.

### Temporal dynamics of the decision circuit when neurons are allowed mixed representations

Next, we examined the temporal dynamics of the decision circuit across layers. To extend our previous population activity analyses (Livi et al., 2025), we constructed soft-assigned profiles in which each neuron contributed to each variable proportionally to its β-weights. Neurons were not assigned to a single variable; instead, weights were derived from normalized β-coefficients, with sign preserved to combine positively and negatively encoding populations.

For each variable class, population activity profiles were computed as weighted sums of all neurons’ ΔF/F traces, a strategy that does not enforce any particular limits on the number or magnitude of encoded variables.

For each variable and layer, trials were divided into quantiles –two for *position of A* and *chosen side*, and three for the remaining variables. ΔF/F traces were aligned to offer onset and choice, and neurons were weighted by their β-based contributions (see **Methods**). As in our previous work (Livi et al., 2025), we pooled *offer value A and B* into one variable (*offer value A*|*B*) and *offer value ipsilateral and contralateral* into another (*offer value I*|*C*).

**Fig.5** and **Fig.6** illustrate the population activity profiles obtained for L2/3 and L5, respectively. These traces were computed keeping separate positive and negative encoding. Direct comparison with the equivalent profiles obtained with categorical encoding (Livi et al., 2025) indicate high similarity.

To compare the temporal dynamics of different neural signals, we next combined contributions with positive and negative encoding. Population activity profiles for L2/3 (**Fig.7A-C**) and L5 (**Fig.7D-F**) were nearly identical to those obtained under categorical encoding (Livi et al., 2025). Specifically: (i) offer value A|B emerged earliest in L2/3; (ii) offer value I|C emerged earliest in L5; (iii) chosen side and chosen value rose around the choice epoch in both layers; and (iv) temporal delays between L2/3 and L5 matched those previously observed.

For each variable, we computed the SOE. We found that SOE traces (**Fig.7G-H**) and their normalized forms (**Fig.7I**) preserved the temporal ordering observed under categorical encoding (**Fig.7J**). Specifically, after the beginning of the offer presentation, first signals to emerge are *offer value A*|*B* and *position of A*, in L2/3, and *offer value I*|*C*, in L5. Then *chosen side* in L5. Similarly to what observed by Livi et al. (2025), this analysis provides two important insights on the architecture of the choice processing. First, *offer value A*|*B* in L2/3 emerges before *offer value I*|*C* in L5 (Sum of Squared Difference [SSD]: 462 ms, Half-Peak Timing [HPT]: 307 ms), suggesting that the initial representation of the offer is in a value space. Second, for both *offer value I*|*C* and *chosen side* the representation emerges first in L5 and later in L2/3 (for *offer value I*|*C*,SSD: 406 ms, HPT: 94 ms; for *chosen side*, SSD: 266 ms, HPT: 48). This temporal dynamics suggests the winner-take-all competition takes place within L5.

Taken together, these results show that allowing neurons to contribute to multiple variables did not alter the population-level temporal structure.

### Functional connectivity of the decision circuit

Last, we focused on GCA. In previous work, we used GCA to assess the functional connectivity of the decision circuit, revealing structured patterns of directed interactions among cell groups defined by encoded variables (Livi et al., 2025). Specifically, we observed an overall flow of functional connections (i.e., *Net Connection Density*) from *position of A* and *offer value A*|*B* cells toward *offer value I*|*C, chosen value*, and *chosen side* cells. Here, we extended this analysis by allowing individual neurons to represent linear combinations of decision variables.

First, we performed GCA on imaging data from each session. As in (Livi et al., 2025), ΔF/F traces were concatenated across trials and modeled using vector autoregressive processes, with model order selected via AIC (Barnett and Seth, 2014). Significant directed influences (p < 0.01, false-discovery-rate corrected) yielded a binarized adjacency matrix per session (see Methods), resulting in a functional connectivity matrix (**Fig.8A, left**).

Second, each significant connection between neurons *n* and *m* was distributed across variable pairs using the outer product of their normalized β-vectors from the LASSO analysis (**Fig.8A, right**). Third, results were aggregated across cell pairs and sessions. Following previous analyses, we computed total Numbers of Connections (**Fig.8B**), Connection Densities (**Fig.8C,D**), and Net Connection Density per variable (**Fig.8E**).

Net Connection Density was computed by summing the densities of received connections (*sink*) and subtracting the densities of sent connections (*source*) (**Fig.8E, upper row**). We also generated bootstrap null distributions and computed the percentile position of the observed net densities relative to the null (**Fig.8E, lower row**).

Two aspects of the results were particularly noteworthy. First, task-related neurons exhibited higher connection densities than expected under the bootstrap null, whereas untuned cells exhibited lower-than-expected densities. This effect is evident when comparing the untuned→untuned entry in **Fig.8C** (density = 5.19) with the tuned→tuned entry (density = 11.88). A similar but weaker pattern was present in the categorical GCA (Livi et al., 2025). Second, the net directional flow of connections remained consistent with earlier work. **Fig.8F** shows the net connections from (Livi et al., 2025), and each variable preserved the directionality of connections, with the exception of untuned cells.

Comparing the present results (**Fig.8B–E**) with those obtained under categorical encoding revealed only modest quantitative differences, primarily affecting untuned versus tuned cells. Specifically, untuned→untuned connections increased in number (not shown), although their relative density decreased slightly, while tuned→tuned connection density increased substantially under soft GCA. We interpret this effect as a consequence of LASSO penalization increasing the number of weakly selective neurons. Importantly, the global circuit structure remained unchanged, reinforcing the conclusion that OFC organization is robustly organized by task variables and does not depend on enforcing a one-cell – one-variable assumption.

In summary, relaxing the one variable constraint did not substantially alter the functional connectivity of the decision circuit.

## Discussion

The present study re-examined the architecture of mouse OFC during economic choice by relaxing the single-variable constraint imposed in our previous analyses (Livi et al., 2025). In that work, we described a laminarly organized decision circuit using a winner-take-all classification scheme. Here, we extend those analyses by applying LASSO regression, allowing individual neurons to encode multiple decision variables.

LASSO regression provided a substantive test of generalizability: the neuronal population was approximately evenly split between neurons best explained by a single variable (categorical) and those exhibiting mixed selectivity. Importantly, these proportions were similar across superficial (L2/3) and deep (L5) layers, suggesting that the encoding scheme is distributed comparably across cortical depth.

Crucially, we caution against over-interpreting these results. Neuronal signals are noisy, the number of trials is limited, and decision variables in OFC are inherently correlated. Under these conditions, it would be premature to draw broad conclusions about the fundamental nature of single-neuron encoding in OFC. These issues are being addressed in ongoing work in non-human primate OFC (Crimmins et al., 2025).

Relaxing the restriction on the number of encoded variables did not alter the main population-level findings in three critical domains: laminar organization, functional connectivity, and temporal dynamics. The same laminar organization of decision signals emerged when neurons were allowed to express linear combinations of multiple variables, indicating that the previously described properties of the decision circuit are not artifactually imposed by the analytical framework, but instead reflect a robust property of the OFC population. Likewise, when functional connectivity (measured via GCA), population activity profiles, and decoding analyses were recomputed using continuous regression weights rather than discrete neuron classes, the resulting temporal structure and interaction patterns closely matched those obtained under the single-variable constraint. This convergence indicates that the population-level functional organization is stable regardless of whether neurons exhibit categorical or mixed selectivity.

Together, these results support a structured and laminar organization of decision-related signals in mouse OFC. Because decision-related variables are encoded later and more robustly in deeper cortical layers, the findings are consistent with the view that decisions emerge, at least in part, from processing within OFC. At the same time, the diversity of single-neuron responses suggests that additional organizational principles may remain to be uncovered. Combining large-scale imaging with targeted perturbations will be essential for determining how neurons across layers, and representing different variables or mixtures thereof, contribute to computation within the OFC circuit.

## Acknowledgements

We thank Timothy Crimmins and Gaia Tavoni for discussions and comments on the manuscript, and for their work on mixed selectivity in orbitofrontal cortex of non-human primate. Their work has been instrumental to the interpretation of the results of this manuscript. This work was supported by the National Institutes of Health (grant numbers R21-DA042882 to C.P.S., R01-DA055709 to C.P.S., R01-DC020034 to T.E.H.) and the McDonnell Center for System Neuroscience (Small Grant to A.L.).

## Notes

**Conflict of interest:** Declared none

### Competing Interest Statement

The authors have declared no competing interest.

